# Gene-tree reconciliation with MUL-trees to resolve polyploidy events

**DOI:** 10.1101/058149

**Authors:** Gregg W.C. Thomas, S. Hussain Ather, Matthew W. Hahn

## Abstract

Polyploidy can have a huge impact on the evolution of species, and it is a common occurrence, especially in plants. The two types of polyploids - autopolyploids and allopolyploids - differ in the level of divergence between the genes that are brought together in the new polyploid lineage. Because allopolyploids are formed via hybridization, the homoeologous copies of genes within them are at least as divergent as orthologs in the parental species that came together to form them. This means that common methods for estimating the parental lineages of allopolyploidy events are not accurate, and can lead to incorrect inferences about the number of gene duplications and losses. Here, we have adapted an algorithm for topology-based gene-tree reconciliation to work with multi-labeled trees (MUL-trees). By definition, MUL-trees have some tips with identical labels, which makes them a natural representation of the genomes of polyploids. Using this new reconciliation algorithm we can: accurately place allopolyploidy events on a phylogeny, identify the parental lineages that hybridized to form allopolyploids, distinguish between allo-, auto-, and (in most cases) no polyploidy, and correctly count the number of duplications and losses in a set of gene trees. We validate our method using gene trees simulated with and without polyploidy, and revisit the history of polyploidy in data from the clades including both baker’s yeast and bread wheat. Our re-analysis of the yeast data confirms the allopolyploid origin and parental lineages previously identified for this group. The method presented here should find wide use in the growing number of genomes from species with a history of polyploidy.

---

Polyploidy as a result of whole genome duplication (WGD) can be a key evolutionary event. At least two ancient WGDs have been postulated at the origin of vertebrate animals (Ohno 1970; Dehal and Boore 2005), with yet another occurring before the radiation of bony fishes (Amores et al. 1998; Van de Peer 2004). Polyploidy events are far more common in plants. It is estimated that approximately 25% of extant angiosperms have experienced a recent polyploidization, while 70% of angiosperm species show signs of a more ancient event (Barker et al. 2015). In fact, it is estimated that between 15 and 30% of speciation in angiosperms results from polyploidy (Wood et al. 2009; Mayrose et al. 2011). The prevalence of these events combined with the success of flowering plants suggests that WGDs must confer some advantage, possibly by increasing speciation rates (Werth and Windham 1991; Lynch and Force 2000; but see Mayrose et al. 2011; Muir and Hahn 2015), by decreasing extinction rates (Crow and Wagner 2006), or by providing species with a large amount of genetic material from which novel phenotypes can arise (Adams and Wendel 2005; Soltis and Soltis 2009; Edger et al. 2015).

Because of their importance in adaptation and speciation, multiple methods have been employed to study polyploidy events. The goals of these methods vary and can include: identifying polyploidy events, placing events in a phylogenetic context to identify the lineages on which they took place, or counting gene duplications and losses in the presence of polyploidy (Table 1). When placing polyploidy events in a phylogenetic context, either relatively or absolutely in time, care must be taken to distinguish between the two types of polyploidy. Autopolyploidy occurs when an individual inherits sets of chromosomes from parents of the same species. Genes duplicated as the result of autopolyploidy are paralogous. Following Glover et al. (2016), we will refer to these as ohnologs, although this term was originally used to describe a gene arising from any type of WGD event (Wolfe 2000). Allopolyploidy occurs when an individual inherits sets of chromosomes from parents of different species through hybridization. Genes duplicated as the result of allopolyploidy are called homoeologs. The term “homoeolog” was originally applied to relationships between chromosomes in allopolyploids (Glover et al. 2016), and seems most appropriate as a descriptor of the genealogical relationships of homologous genes within allopolyploids: not quite orthologs and not quite paralogs (Glover et al. 2016). Recognizing that homoeologs have relationships differing from those occurring between homologous genes within autopolyploids is key in preventing the mis-classification of allopolyploids as autopolyploids (cf. Doyle and Egan 2010).

**Table 1.**
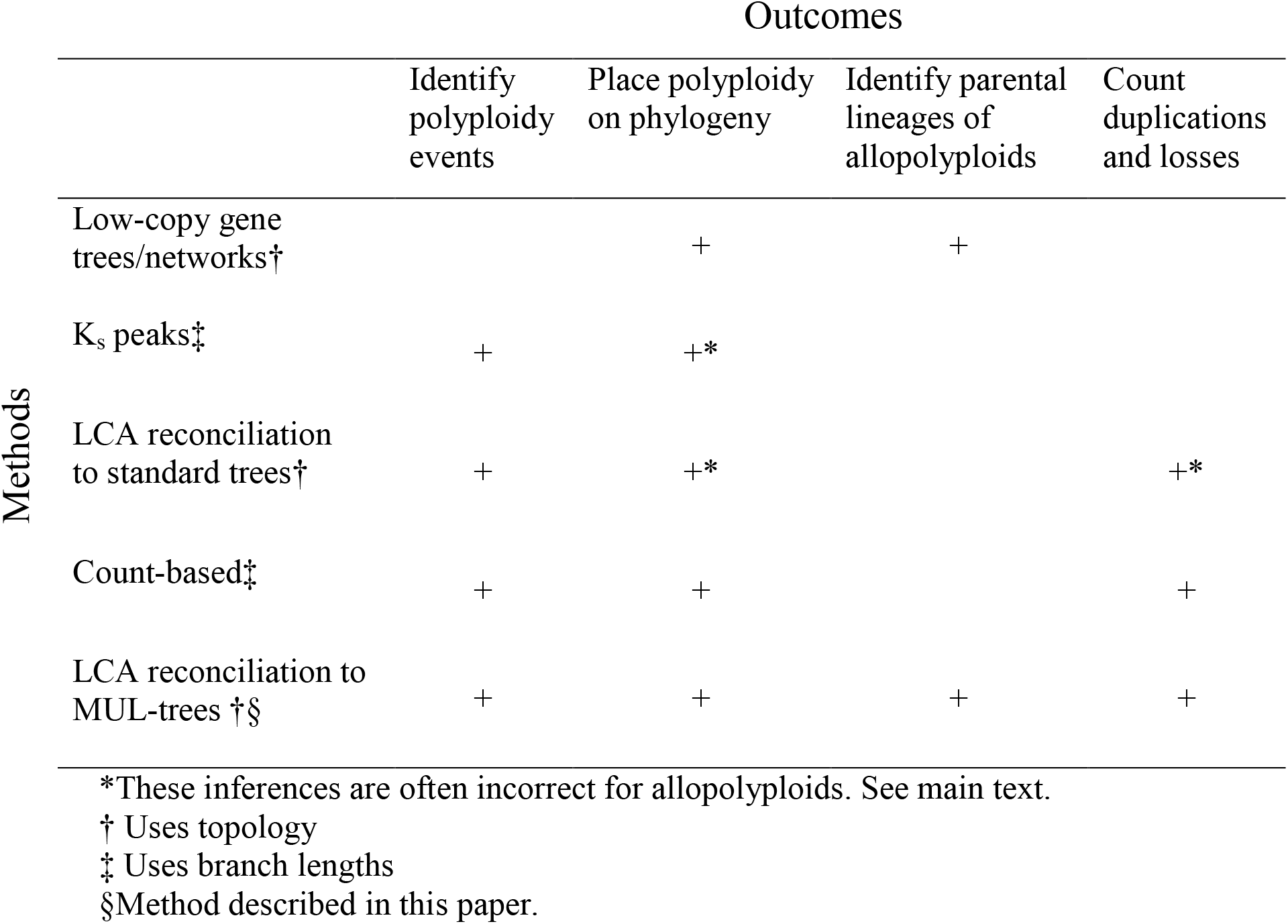

There are many different methods used to identify and study polyploidy events. Non-phylogenetic methods, such as synteny and karyotyping, have long been used to identify polyploid species and will not be discussed further here (for review see Barker et al. 2015; Kellogg 2016). Phylogenetic methods (Table 1) use the relationships of genes between closely related species to identify and place polyploidy events on a phylogeny. One common approach is to build phylogenies of low-copy genes and to treat the resulting gene trees as species relationships (see Table S1 for a summary of studies that have used each method). Because these are low-copy genes, only polyploid species should be represented with multiple copies, in which case each copy represents either an ohnolog or a homoeolog. If the polyploid species or clade in question is an autopolyploid, the ohnologs will be sister to each other in the genome tree (topology in Fig. 1a); if they are allopolyploids, the homoeologs will be sister to different diploid taxa (as long as these lineages are sampled; topology in Fig 1b). These diploid taxa are either the direct progenitors of the allopolyploid species (the parental *species)* or the extant taxa most closely related to the progenitors (the parental *lineages*; Fig S1) if the direct progenitors are extinct or were not sampled. Such methods can benefit from including more species closely related to the parental lineages of the polyploids in order to separate multiple WGDs close in time. Incongruence caused by error in gene tree inference or incomplete lineage sorting can be addressed by sequencing more genes (e.g. Brassac and Blattner 2015), or by comparing nuclear loci to genes in the mitochondria or chloroplast (e.g. Popp and Oxelman 2001), and methods have been devised to make consensus phylogenies from sets of gene trees (e.g. Huber et al. 2006; Marcussen et al. 2012; Marcussen et al. 2015). However, the use of low-copy genes is only applicable for more recent polyploidy events, as additional paralogs created by gene duplication (and the loss of homoeologs) are not considered by these methods (Marcussen et al. 2012; Jones et al. 2013).

**Figure 1.**
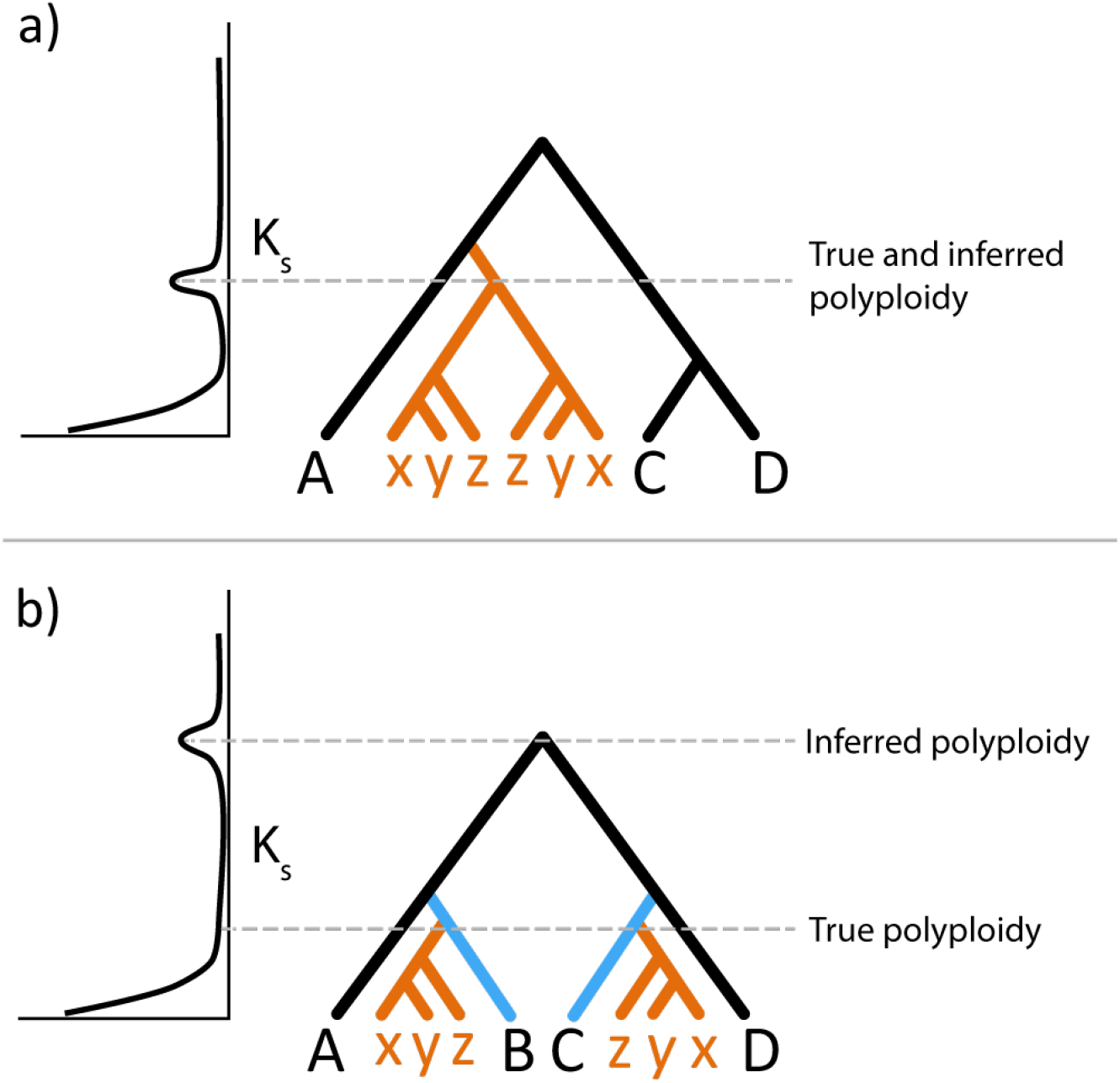
Methods to identify the placement of WGD in a phylogeny. (***a***) K_s_-based methods can correctly date cases of autopolyploidy. Here the peak in the distribution of K_s_ values between paralogs in a single species (shown as a density plot on the left) corresponds to the duplication node in the tree. (***b***) K_s_-based methods incorrectly date cases of allopolyploidy. Here a hybridization event occurred between species B and C (in blue) resulting in the allopolyploid lineage that gave rise to the XYZ clade (orange). In cases like this the peak in the distribution of K_s_ for a single species corresponds to the most recent common ancestor of the two parental lineages, rather than the timing of the WGD. Similar results would be found using LCA reconciliation methods discussed in the text.

In order to identify previously unknown polyploidy events, K_s_-based, least common ancestor (LCA) reconciliation, and count-based methods are used (Table 1; Table S1). K_s_-based methods start by identifying pairs of duplicate genes within a species of interest, then measuring the synonymous divergence, or K_s_, between them. In a species that has not experienced polyploidy, the expectation is that most duplicates will be very recent - and have a low K_s_ - while very few pairs will have high K_s_ (Lynch and Conery 2000). However, in a lineage in which polyploidy has occurred, peaks observed in this distribution are said to correspond to a burst of duplications from the WGD, though this is not always the case (see below; Lynch and Conery 2000; Blanc and Wolfe 2004). Peaks observed in a single species are placed on the tip branch of a species phylogeny, while peaks of K_s_ shared among multiple species are placed on internal branches (e.g. Cui et al. 2006; Barker et al. 2008; Barker et al. 2009).

LCA-based methods consider the topology of gene trees, and attempt to map duplication events onto an accepted species phylogeny via gene tree-species tree reconciliation (Goodman et al 1979; Page 1994). Gene duplications on each branch of a species tree can be counted, and branches with an unusually large number of gene duplications per unit time can indicate a polyploidy event (Cannon et al. 2015). Often LCA reconciliation is carried out on simplified gene trees (Bowers et al. 2003; Li et al. 2015), but the inferences remain the same. Finally, recent likelihood-based “count” methods use gene copy-numbers to identify branches with more gene duplications than expected without polyploidy (Rabier et al. 2014; Tiley et al. 2016). Neither gene trees nor pairwise distances are calculated among homologs, and instead the number of gene copies in each genome is used as a character that evolves across a species phylogeny. K_s_, LCA, and count-based methods can be used for analysis of polyploidy events at any depth in time, though they are most commonly used for more ancient events where one or both parental lineages are extinct.

However, the K_s_ and LCA methods described above do not give a full picture of the evolutionary history of a polyploidy event, and may be positively misleading about multiple aspects of WGDs. This is because these methods do not take into account the difference in the degree of divergence between duplicate gene pairs in the two types of polyploidy. Specifically, K_s_- and LCA-based methods fail because they treat the two homologous copies of a gene arising from polyploidy as paralogs regardless of their mode of origin - that is, homologous genes related by a duplication event at their most recent common ancestor (Fitch 1970). However, for allopolyploids, these two genes are more akin to orthologs, as they are actually related by a speciation event at their most recent common ancestor (Glover et al. 2016). While these methods will work for autopolyploids—since the divergence time of ohnologs is the true timing of the WGD (Fig. 1a)—for allopolyploids they incorrectly identify the most recent common ancestor of the species that hybridized to form the polyploid as the time that the WGD occurred (Fig. 1b) (Doyle and Egan 2010; Kellogg 2016). When using LCA-based gene tree methods, WGDs due to allopolyploidy can also lead to incorrect inferences of many additional gene duplications and losses when none have occurred (see Methods).

One major issue shared by K_s_-based, LCA, and gene count methods when used to study polyploidy is that the typical bifurcating, singly-labeled representations of species relationships (henceforth referred to as singly-labeled trees) only represent one of the multiple homoeologous sub-genomes (sets of chromosomes) in any allopolyploid species (Fig. 2a). Species networks are the correct representation for allopolyploids as they can highlight both the parental lineages involved and the relative timing of the hybridization event (Fig. 2b) (Linder and Rieseberg 2004; Huber and Moultan 2006; Jones et al. 2013). However, networks represent species relationships, and therefore can be less practically useful for analyses involving multiple individual genes in allopolyploid genomes. An alternative and fully equivalent representation to the species relationships in networks uses multi-labeled trees (MUL-trees, sometimes referred to as “genome trees” in this context) to represent genome relationships for polyploidy events (Fig. 2c) (Huber et al 2006; Lott et al. 2009). MUL-trees are trees in which the tip labels are not necessarily unique (Huson et al 2006); this allows one to represent all sub-genomes in an allopolyploid as descendants of different parental lineages (Fig. 2c), or as descendants of the same lineage for autopolyploids (e.g. Fig. 1a).

**Figure 2.**
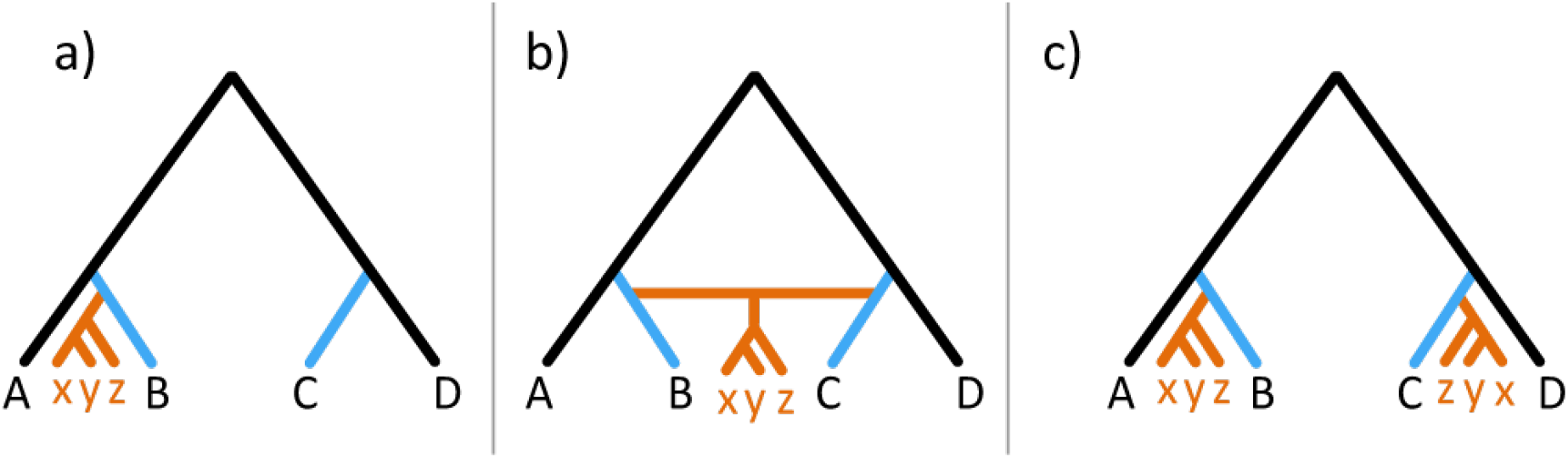
Representation of allopolyploid clades. Given that a hybridization event occurred between species B and C (in blue) resulting in the allopolyploid lineage that gave rise to the XYZ clade (orange), species relationships can be represented in three ways. (***a***) Most species tree reconstruction methods represent each species with a single label. This is not correct for polyploid species as these singly-labeled species trees incorrectly represent each polyploid species with a single label, showing only one of the two possible topologies. (***b***) Phylogenetic networks correctly display the species-level phylogeny by showing hybridization events. (***c***) Multi-labeled trees (MUL-trees) are genome-level phylogenies equivalent to networks, and therefore both sub-genomes of the polyploid species are represented and it is easy to identify the parental lineages involved in the hybridization event. The relative placement of hybridization in the phylogeny is implicit in this representation as the point at which both sub-genomes originate.

Here, we have adapted the LCA algorithm for use with MUL-trees, implementing this method in the software package GRAMPA (Gene-tree Reconciliation Algorithm with MUL-trees for Polyploid Analysis). This representation and algorithm allows us to correctly infer gene duplications and losses in the presence of polyploidy, and to identify the most likely placement of polyploid clades and their parental lineages. In most cases, it should also be able to infer whether or not a polyploidy event has taken place. We demonstrate that this new method works on simulated data, and we revisit two different datasets that include allopolyploid species, confirming a newly presented conclusion on the parental lineages leading to the clade that includes baker’s yeast.

## Methods

### Algorithm

While the problem of reconciling gene trees to reticulated phylogenies has been explored before (Yu et al. 2013; To and Scornavacca 2015), we have devised an LCA mapping algorithm that reconciles gene phylogenies to genome relationships represented as MUL-trees, a natural representation of polyploidy events. The LCA mapping algorithm is a method that identifies and counts duplication and loss events on a gene tree given an accepted singly-labeled species tree (Goodman et al 1979; Page 1994). It can also be used for species tree inference by searching for the species tree that minimizes the total number of duplications and losses inferred given a set of gene trees (Guigó et al. 1996). The main hurdle in applying LCA mapping to MUL-trees is that, when reconciling to a MUL-tree, some nodes have more than one possible map. In particular, some tip nodes cannot be initialized with a single map (because tips are necessarily not uniquely labeled), which subsequently allows internal nodes to also have more than one possible map. We side-step this problem by trying all possible combinations of initial tip maps and applying the parsimony assumption – that the correct map will have the lowest score.

In standard LCA mapping each internal node in the gene tree, *n_g_*, is defined by the set of species at the tips below it in the tree. The same node is also associated with a node in the species tree, *n_s_*, through a map, *M*(*n_g_*). *M*(*n_g_*) = *n_s_*, where *n_s_* is the node in the species tree that is the least common ancestor of the species that define *n_g_*. For example, in the gene tree depicted in Figure 3a, node 1G is defined by the tips A1 and B1. These tips map to species A and B, respectively, and the first node in the singly-labeled species tree (going from tips to root) that includes species A and B is 1S (Fig. 3b). Therefore, *M*(1*G*) = 15. This process is repeated for every internal node in the gene tree until all nodes are mapped. For each *n_g_* there is a single possible node in the species tree to which it can map; however, nodes in the species tree can have multiple nodes map to them. Nodes in the gene tree are said to be duplication nodes when they map to the same species tree node as at least one of their descendants (for example, nodes 3G and 4G in Fig. 3b).

**Figure 3.**
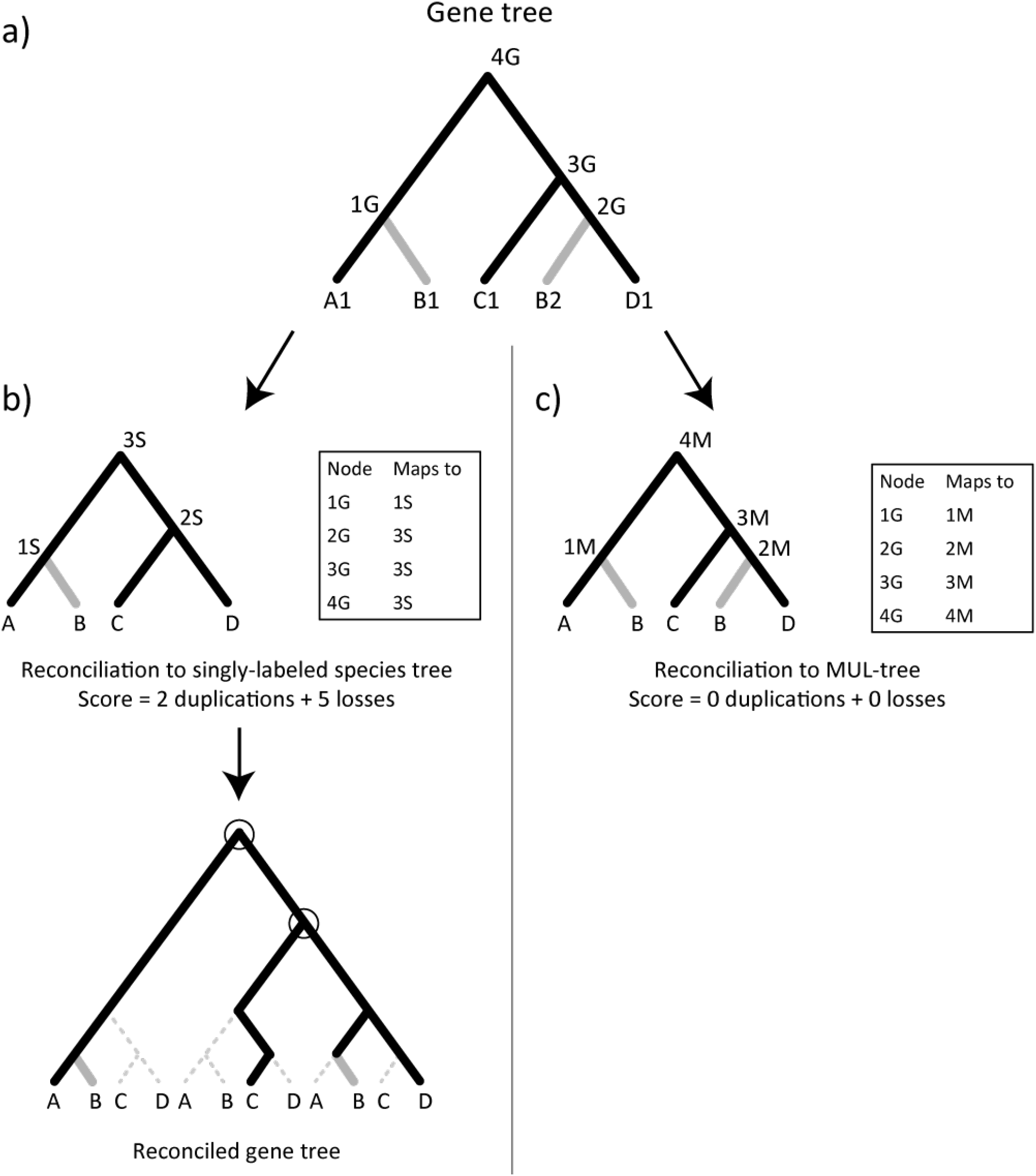
Reconciliation with a gene tree from an allopolyploid lineage. (***a***) A representative gene tree is shown, with every homolog labeled, including the two homoeologs from allopolyploid species B. Internal nodes are also labeled to understand the mappings below. (***b***) Top: Reconciliation to a pre-defined singly-labeled species tree, with maps between gene tree nodes and species tree nodes shown (species tree nodes are labeled 1S, 2S, and 3S). In this case, genes B1 and B2 are treated as paralogs and extra duplications and losses are inferred, and the duplications are placed ancestral to the actual parental lineages (A and D) of the allopolyploid. Bottom: The reconciled gene tree showing the duplications (circles) and losses (dashed branches) inferred. (***c***) Reconciliation to a pre-defined MUL-tree, with both allopolyploid sub-genomes represented, with maps between gene tree nodes and MUL-tree nodes shown (MUL-tree nodes are labeled 1M, 2M, 3M, and 4M). In this case, genes B1 and B2 are treated as homoeologs and using our algorithm no extra duplications or losses are inferred.

When the mapping of a gene tree (*T_G_*) is performed, nodes in the gene tree are classified as either duplication or speciation nodes, based on the procedure described above. The number of duplications that occur in the gene tree (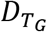) is then just the number of duplication nodes. Losses along a branch in the gene tree, *b*_g_, subtended by *n_g_* are counted as the difference in the depth of the maps of a node in the gene tree and its ancestor (Durand et al. 2006):

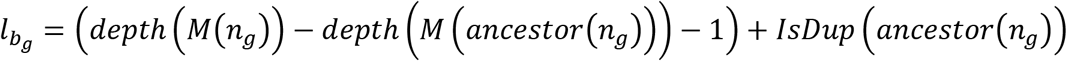

Where *IsDup* is a function that returns 1 if the ancestor of *n_g_* is a duplication node and 0 otherwise. The depth of a node refers to its distance from the root in the species tree. The node at the root has a depth of 1, with each node farther down the tree having depth +1 from its ancestor. For example, in Fig. 3b, node 3S (the root) has a depth of 1, node 1S has a depth of 2, and nodes A and B each have a depth of 3. The total number of losses that occurred on the gene tree (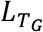) is simply the sum of the number of losses on all branches, 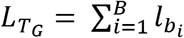., and the reconciliation score (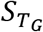) for this gene tree is:

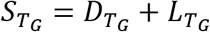

Many reconciliation algorithms do not count losses in cases where the root of a gene tree does not map to the root of the species tree. This happens because there are no nodes in the gene tree mapping to entire branches of the species tree, and therefore no calculations can be performed. In these cases (when *n_g_* = root) we add *depth*(*M*(*n_g_*) — 1) to 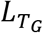 because the most parsimonious solution is simply a single loss of each branch above this node in the species tree.

The entire reconciliation process hinges on the fact that the mapping function is initialized with the tips of the gene tree mapped to their corresponding species label in the species tree. In a MUL-tree, repeated clades represent the sub-genomes of the polyploid species, and their placement in the MUL-tree defines parental lineages of the polyploid event (Fig. 2c, Fig S1). Given a gene tree, we proceed with the LCA mapping algorithm as described above, except that any tip that maps to a polyploid species now has two possible initial maps: to either of the sub-genomes represented in the MUL-tree (species “B” in Fig. 3c). This leads to unresolved internal maps and the inability to classify nodes correctly. To solve this problem, we run LCA mapping with a tip within a polyploid clade initialized to one sub-genome first, and then we run LCA mapping again with that same tip initialized to the other sub-genome, giving us two maps and two reconciliation scores for the single gene tree. We then apply the parsimony principle for these two possible maps: whichever initial mapping results in the lowest score is the correct map. If there is more than one gene in the gene tree from the polyploid clade we try all possible combinations of initial maps. In cases where multiple different mappings are all tied for the lowest score, we report all possible mappings.

The MUL-tree reconciliation algorithm is applicable for any number of genes in any number of polyploid species, but the algorithm becomes very slow for large polyploid clades because of the large number of combinations of initial maps to consider. Given that a gene tree has *m* genes represented from polyploid species, the run time for mapping this gene tree is *o*(2*^m^n*), since the mapping algorithm itself is linear for a tree with *n* nodes (Zmasek and Eddy 2001) and we perform 2*^m^* maps (Fig. S2b). A similar brute force method was devised by Yu et al. (2013) when mapping alleles in a gene tree to a species tree in order to infer hybridization. In the context of polyploidy, we devised several methods to expedite the process of choosing the correct map by using context within both the gene tree and the MUL-tree. If any genes from different polyploid species form a clade within the gene tree, then the most parsimonious solution will always have them mapping to the same sub-genome in the MUL-tree. We group these nodes together, essentially treating the clade as a single tip of the tree, and try both maps on the group as a whole rather than on each tip individually (Fig. S2c). We also consider the species sister to the polyploid clades in the MUL-tree. If we observe clades in the gene tree that include only polyploid species and these sister species, then the most parsimonious initial map for that group is guaranteed to be the one in the MUL-tree with the corresponding sister species. The algorithm therefore fixes the maps of these nodes before proceeding (Fig. S2d). This method gives a reconciliation score for mapping a single gene tree to a single MUL-tree, and can be applied to a set of gene trees to obtain a total reconciliation score for a MUL-tree. Even with the speed-ups described above, some gene trees can still take an exorbitant amount of time to reconcile. We therefore place a cap of 15 on the number of groups that can be considered for a given gene tree. This limits any gene tree to at most 2^15^ = 32,768 possible maps. Gene trees over this cap are skipped. The number of groups in a gene tree can vary for different MUL-trees. To ensure consistency when comparing scores between MUL-trees, we first calculate the number of groups for each gene tree/MUL-tree combination and filter out gene trees that are over the cap for any MUL-tree.

Thus far we have assumed that the placement of the polyploidy event is already known, since a single MUL-tree represents a single polyploid scenario. When it is not known, we have implemented a search strategy to find the most parsimonious placement of a polyploidy event, given a singly-labeled species tree. We define two nodes of interest in a singly-labeled species tree that we use to build a MUL-tree. Node H1 defines the location of the sub-genome for the polyploid species represented in the singly-labeled species tree (as in Fig. S3a). Node H2 defines the location of the second parental lineage and unrepresented polyploid sub-genome (Fig. S3a). When H1 is specified, the sub-tree that is rooted by it and the branch that it subtends are copied and placed on the branch that is subtended by H2. Our modified LCA mapping algorithm is then run on the resulting MUL-tree, and a total reconciliation score is obtained by summing across scores for all gene trees. The algorithm can be limited to a specified pair of H1 and H2 nodes, or only a specified H1 node (searching for H2), or no nodes specified (searching for both H1 and H2). The MUL-tree defined by a particular H1 and H2 with the lowest total reconciliation score reveals the location and type of the most parsimonious polyploidy event. Placement of H1 and H2 as sister to different lineages indicates allopolyploidy, while placement of H1 and H2 on the same node in the species tree represents an autopolyploid event. We also perform LCA reconciliation to the input singly-labeled tree. In many instances, if no polyploidy has occurred in the sampled lineages, the singly-labeled tree will return the lowest score.

GRAMPA’s search strategy guarantees that it will be able to distinguish between allo-, auto-, and no polyploidy in most cases (see Results). However, there are circumstances in which the complete extinction of both parental lineages of an allopolyploid would lead us to incorrectly infer autopolyploidy. The distinction between parental lineages and parental species is important when discussing allopolyploids. We define a parental lineage of an allopolyploid as the clade of species more closely related to one sub-genome than the other (Fig. S1). A parental lineage can include the direct parental species that hybridized to form the allopolyploid (if they are extant), but also can include species closely related to the parents. For example, in Fig S1a, the true network shows that the species A' and B' hybridized to form the polyploid lineage P. If both parental species are extant and sampled, it is clear that we will recover the proper placement of the hybridization event (Fig S1a). However, both of these parental species also have sister species (A and B), which are part of the two parental lineages. Even with extinction of one (Fig S1b) or both parental species, our method will still be able to identify the parental lineages, and therefore correctly infer allopolyploidy. These two cases (Figs. S1a and b) are sometimes referred to as “neopolyploidy” (Mandakova et al. 2010). Similarly, in the case of the complete extinction of one parental lineage (Fig S1c; referred to as “mesopolyploidy”), our method will still implicitly identify such a lineage via the placement of H1 or H2 on an internal node of the singly-labeled tree. The only instance in which our method leads to an incorrect inference of the type of polyploidy is in the case of the complete extinction of both parental lineages (Fig S1d; sometimes referred to as “paleopolyploidy”), in which case autopolyploidy will be inferred. These definitions of neo-, meso-, and paleopolyploidy are based on genealogical context alone and differ from those based on cytology (e.g. Mandakova et al. 2010).

### Simulations

We first checked that our modified LCA mapping algorithm counted the correct number of duplications and losses by manually reconciling a small set of 25 gene trees onto 8 MUL-trees and 1 singly-labeled tree to represent varying cases of gain, loss, and polyploidy. We generated our MUL-trees by starting with a single arbitrary singly-labeled species tree topology (Fig. S3a) and specifying a node as H1. With that node as H1 we tried every possible placement of node H2 to construct the MUL-trees (one example is shown in Fig S3b). Gene trees were made by randomly adding or removing branches from these nine trees. Our method always agreed with the expected counts for each type of event (Table S2).

Next we used gene trees simulated in the GuestTreeGen program within JPrIME (Sjöstrand et al. 2013) with varying levels of gene gain and loss and incomplete lineage sorting (ILS) to test the search feature of our algorithm. Specifically, we want to know if, given sets of gene trees simulated under conditions of allo-, auto-, or no polyploidy, our algorithm correctly identifies the type of polyploidy that has occurred and the parental lineages involved in the polyploidization event (Fig S4). To simulate scenarios with polyploidy, we started with an arbitrary MUL-tree with one clade represented twice, indicating both sub-genomes of a set of polyploid species (Fig S4a and b). JPrIME does not accept MUL-trees as input, so we added temporary marker labels to species within one of the polyploid clades. JPrIME then generated 1000 gene trees with this labeled MUL-tree as input. Gene trees were simulated under five scenarios of gain and loss rates and three scenarios of ILS, giving us 15 sets of 1000 gene trees each for each starting tree of allopolyploidy (Fig S4a), autopolyploidy (Fig S4b), and no polyploidy (Fig S4c). We removed the marker labels from the gene trees and used them as input for our algorithm, along with a singly-labeled tree in which only one of the polyploid clades is represented. We then searched for the H1 and H2 nodes that minimized the reconciliation score. For the simulations of allopolyploidy, we also gave as input to our algorithm in a separate run the alternate singly-labeled topology with the other polyploid clade represented. In all, this resulted in 45 simulated datasets and 60 inferences by our algorithm.

### Yeast

The yeast data came from the study of Marcet-Houben and Gabaldón (2015). We use the species tree topology depicted in their supplementary figures with all 27 species present (Fig. S5). We downloaded the set of 5,402 gene trees used by Marcet-Houben and Gabaldón (2015) from PhylomeDB (Phylome ID: 206). These were the main inputs to our algorithm, along with the baker’s yeast clade set as the H1 node (node n5 in Fig. 4a and Fig. S5). We then let our algorithm search for the optimal placement of H2, which would allow for identification of parental lineages of this polyploidy event.

**Figure 4.**
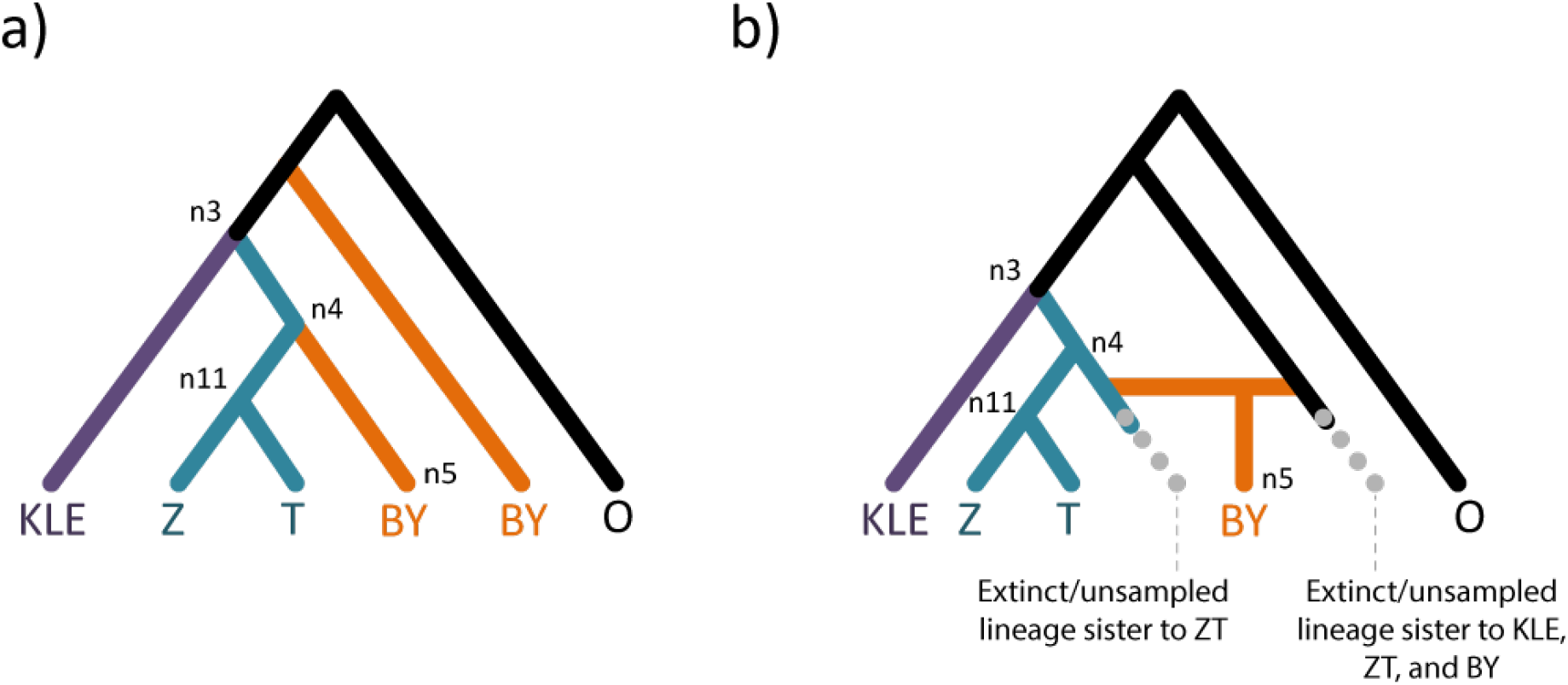
The inferred MUL-tree and species network for the baker’s yeast data. BY: baker’s yeast clade, Z: *Z*. *rouxii*, T: *T*. *delbruekii*, KLE: KLE clade, O: Outgroups. See Supplementary Figure S3 for full tree and all node labels. (***a***) The optimal MUL-tree inferred when searching for both H1 and H2. H1 was confirmed to be on the branch leading to the baker’s yeast clade (as normally represented in a species tree; specified by node n5), while H2 was inferred to be ancestral to the KLE, ZT, and BY clades (n3). (***b***) The optimal MUL-tree as a network. This representation highlights the two parental lineages of the allopolyploid event: an extinct or unsampled lineage sister to the ZT clade, and an extinct or unsampled lineage sister the KLE, ZY, and BY clades.

Several steps were required to prepare the gene trees for our program. 779 trees consisted of labels that were either identical to or a subset of another tree. These 779 trees were removed. Notung version 2.8 (Durand et al. 2006) was used to root the gene trees and to perform bootstrap rearrangement with a 0.7 bootstrap threshold. Bootstrap rearrangement ensures more accurate gene trees by finding the most parsimonious (with respect to duplications and losses) topology around nodes with low bootstrap support by finding the lowest scoring topological ordering of affected taxa. Finally, we capped our algorithm to consider at most 10 collapsed groups (Fig. S1). This step cut out an additional 636 trees leaving us with a final set of 3,987 gene trees to reconcile. We followed a similar procedure for the set of 963 trees containing “ohnologs” obtained from Marcet-Houben and Gabaldón (2015) (http://genome.crg.es/~mmarcet/yeast_hybrids/phylome_table.htm), resulting in 505 usable gene trees. Analysis of these trees containing homologs believed to be due to WGD yielded identical results as those of the whole dataset.

### Wheat

We downloaded 55,519 gene trees from Ensembl Plants v31 (ftp://ftp.ensemblgenomes.org/pub/plants/release-31/emf/ensembl-compara/homologies/) (Kersey et al. 2016). We also obtained the species tree that Ensembl uses for its analyses (Kersey et al. 2016). Both gene trees and species trees were pruned using Newick Utilities (Junier and Zdobnov 2010) so that they only included 10 Poaceae species, including the hexaploid bread wheat species, *Triticum aestivum* (Fig. S6). These species, along with all three bread wheat sub-genomes, are depicted in Supplementary Figure S6. Because the D sub-genome of *Triticum aestivum* may be the result of homoploid hybridization between the A and B sub-genomes (Marcussen et al. 2014), its placement in the singly-labeled species tree is somewhat arbitrary. We chose to place the D sub-genome and its related species *Aegilops tauschii* as sister to the A sub-genome and its related species, *Triticum urartu,* in our representation of the species tree. The gene trees were filtered so they only contained between 5 and 100 tips. This left 9,147 gene trees on which to run our algorithm.

We proceeded by removing one of the three sub-genomes (A, B, or D) from the species tree and pruning all genes that originated from that same sub-genome from the gene trees. For the remaining genes, we masked all labels that specify their sub-genome of origin. We then gave our algorithm this set of gene trees along with a singly-labeled species tree with only one of the two bread wheat sub-genomes represented. We allowed our algorithm to search for H1 and H2 with the expectation that the optimal MUL-tree will correctly represent both wheat sub-genome:

## Results

### Performance of algorithm

Our algorithm is implemented in the software package GRAMPA (Gene-tree Reconciliation Algorithm with MUL-trees for Polyploid Analysis; available at https://github.com/gwct/grampa). The main inputs of the program are a species tree (singly-labeled or MUL) and a set of gene trees. If a singly-labeled tree is input, possible polyploid lineages may be specified (H1 nodes; Fig. S3) along with possible placements of the second parental lineage (H2 nodes; Fig. S3). With H1 specified, GRAMPA will search for the optimal placement of the H2 node. If no H1 and H2 nodes are defined, GRAMPA will generate MUL-trees based on all possible H1 and H2 nodes. In either case, GRAMPA will return a reconciliation score for each MUL-tree considered (as well as for the original singly-labeled tree), including the total number of duplications and losses for each individual gene tree. If a MUL-tree is input (i.e. H1 and H2 specified), GRAMPA will return a total reconciliation score for the tree and individual duplication and loss scores for the gene trees.

We checked that our modified LCA mapping algorithm counted the correct number of duplications and losses by manually reconciling a small set of 25 gene trees onto 8 MUL-trees that represent varying cases of gain and loss (Table S2). Our method always agrees with the expected counts for each type of event. GRAMPA’s search method was then validated using larger sets of gene trees simulated using JPrIME (Sjöstrand et al. 2013) with varying rates of gene gain and loss, and varying amounts of ILS. In every simulation scenario with polyploidy that we tested GRAMPA returned the expected MUL-tree, indicating that we can correctly distinguish between allo- and autopolyploidy, and that in cases of allopolyploidy we can correctly identify the parental lineages that gave rise to the polyploid species (Table S3). For example, when GRAMPA was given a set of gene trees simulated from an allopolyploidy event (Fig S4a) and a corresponding singly-labeled species tree with only one sub-genome represented, it always found the correct MUL-tree. Similarly, given a set of gene trees simulated from an autopolyploidy event (Fig S3b), GRAMPA always returns the correct autopolyploid MUL-tree. These results remain true regardless of gene gain and loss rates or levels of ILS, though with increasing ILS rates we observe inflated counts of gene duplication and loss, as expected with an LCA-based algorithm (Hahn 2007). We also assessed GRAMPA’s performance when given a singly-labeled species tree (Fig. S4c) and gene trees simulated from that species tree; in this scenario no polyploidy has occurred. Reconciliation to the input singly-labeled species tree correctly resulted in the lowest score, indicating that we are able to identify when no polyploidy has occurred.

Distinguishing between allo-, auto-, and no polyploidy simply based on gene-tree topologies is possible because of the penalties that naturally arise when reconciling to the incorrect species topology. For instance, if allopolyploidy has occurred most gene trees should have two copies of the polyploid species present, inducing a penalty of at least one duplication and one loss when reconciling to a singly-labeled tree with only one copy represented (Fig. S7a). This penalty is increased by one loss with each additional lineage between the two polyploid clades (Fig. S7a, gene tree 1). Asymmetric gene loss between the two homoeologous sub-genomes can also provide a signal of allopolyploidy, with extra duplications and losses inferred for gene trees in which homoeologs are present only in the sub-genome that is not represented in the singly-labeled tree (Fig S7a, gene trees 2 and 3). Additional polyploid species make asymmetric gene loss more easily detectable, increasing power to detect polyploidy. A similar penalty occurs when reconciling gene trees resulting from autopolyploidy to a singly-labeled tree (Fig S7b), though this penalty is less pronounced since there is only one possible singly-labeled tree, making asymmetric gene loss indistinguishable.

### Analysis of baker’s yeast

We revisited the interesting case of the WGD occurring in the ancestor of *Saccharomyces cerevisiae* (baker’s yeast), a well-known example of polyploidy (Wolfe and Shields 1997; Kellis et al. 2004). While early authors were circumspect about whether this WGD was an auto- or allopolyploid (Wolfe and Shields 1997; Kellis et al. 2004), Marcet-Houben and Gabaldón (2015) recently postulated that this clade (labeled as the “BY” clade in Fig. 4) was the result of an allopolyploidy event. These authors detected a mismatch in the timing of duplications inferred by count methods and reconciliation, as would be expected given allopolyploidy. This led to the conclusion that an ancient hybridization occurred to create an allopolyploid. This hybridization was inferred to have been between an ancestor of the ZT clade (Fig. 4) and an extinct lineage sister to the KLE, ZT, and BY clades (node n3 in Fig. 4). However, the phylogenetic methods employed by the authors to identify the parental lineages of the allopolyploid could not naturally deal with reticulation in an allopolyploidy event, and thus may have been misled by problems similar to those outlined in the Introduction.

We used 3,987 gene trees (after filtering, see Methods) across 27 yeast species (Fig. S1) from Marcet-Houben and Gabaldón (2015) to reinvestigate the polyploid history of baker’s yeast using GRAMPA. We observed that the optimal MUL-tree inferred by GRAMPA has a reconciliation score of 144,166. We then compared this score to the scores of MUL-trees representing three alternative hypotheses. First, we wanted to confirm that we could detect the WGD by comparing the reconciliation score of the lowest scoring MUL-tree to that of the singly-labeled tree—this tree had a score of 169,031. The fact that allopolyploidy scored much lower than no polyploidy indicates that enough phylogenetic signal remains in these species to differentiate the two scenarios. This signal is enhanced by the fact that there are 12 polyploid species represented in the gene trees, meaning that asymmetric gene loss between sub-genomes is more likely to be detected. We also compared scenarios of auto- vs. allopolyploidy. The autopolyploid MUL-tree had a score of 179,636, and because the optimal allopolyploid MUL-tree has a score much lower than this, we confirm the result from Marcet-Houben and Gabaldón (2015) that the modern baker’s yeast clade is the result of a hybridization event.

We confirmed that GRAMPA’s optimal MUL-tree also corresponds to the specific allopolyploid scenario proposed by Marcet-Houben and Gabaldón (2015; Fig 4a). These results suggest that the most probable parental lineages are an extinct lineage sister to the clade formed by *Z*. *rouxii* and *T*. *delbrueckii* (the so-called ZT clade) and an extinct lineage sister to the KLE, ZT, and the modern BY clades (Fig. 4). The next lowest scoring MUL-tree had about one thousand more duplications and losses, and would have placed the second parental lineage as an extinct lineage sister to only the ZT and modern BY clades. Our results therefore support the claim of Marcet-Houben and Gabaldón (2015) for an allopolyploid origin of the clade including baker’s yeast, and further confirm their inferred parental lineages of this clade.

### Analysis of bread wheat

We also applied GRAMPA to 9,147 gene trees from the clade including the hexaploid species, *Triticum aestivum,* commonly known as bread wheat. This species is the result of three hybridization events, two leading to a WGD (Petersen et al. 2006) and one resulting in a homoploid hybrid species (Marcussen et al. 2014). Analysis of this clade is an especially useful example to demonstrate the accuracy of GRAMPA because the relationships between sub-genomes are known, and genes have been assigned to their sub-genome of origin. With genes labeled according to sub-genome, standard reconciliation can be performed in an approach similar to ours but with a pre-labeled MUL-tree (Altenhoff et al 2015; Bolser et al. 2015). It also presents an interesting test because the current implementation of our algorithm is only designed to map one WGD per tree.

To show that GRAMPA is able to recover the correct MUL-tree for the clade including *T*. *aestivum* and nine other Poaceae species (Fig. S6), we started by analyzing genes from two sub-genomes at a time and removing all labels associating genes with sub-genomes. When we allowed GRAMPA to search for the optimal MUL-tree, we recovered the one with correct sub-genome relationships every time (Fig. S8, Table S5). We also investigated GRAMPA’s results without removing genes from any of the three sub-genomes. Interestingly, when presented with a singly-labeled tree with only one sub-genome represented (Fig. S9a), GRAMPA’s two lowest scoring MUL-trees were those in which the other two un-represented sub-genomes were identified as the H2 clades (Fig. S9b and c). This behavior is especially useful because it implies that GRAMPA could be used to search for multiple allopolyploidy events.

## Discussion

We have developed a method, GRAMPA, to accurately identify whether a polyploidy event has occurred and to place it in a phylogenetic context. This allows us to identify the parental lineages of the polyploid species resulting from the hybridization in the case of allopolyploidy. Our method also allows us to accurately infer the number of duplications and losses in a clade containing an allopolyploid. Though reconciliation methods on reticulated phylogenies have been explored before (Yu et al. 2013; To and Scornavacca 2015), this is the first general method that we know of that performs these types of analyses in the context of polyploidy and is applicable in a wide variety of contexts. Using our method to re-analyze the neo-allopolyploid bread wheat, the results of our algorithm always align with the accepted relationships between sub-genomes. Application to the WGD in the baker’s yeast clade has also confirmed both an allopolyploid origin and the parental lineages inferred previously (Marcet-Houben and Gabaldón 2015). Our method works because it considers the genes duplicated during polyploidization in all possible genealogical contexts: as paralogs in the case of no polyploidy, as ohnologs in the case of autopolyploidy, and as homoeologs in the case of allopolyploidy. K_s_- and standard LCA-based methods fail because they do not make these distinctions.

There are cases where our method may incorrectly report that no polyploidy has occurred, even when it has. The most likely situation in which this will occur is when so many of the ohnologs or homoeologs from the WGD have been lost that the singly-labeled tree has the lowest reconciliation score. This scenario is challenging for all methods that attempt to identify WGDs (Table 1). For genes without any sort of homolog in the same genome, reconciliation to MUL-trees requires a gene loss. Because of this cost, it may seem intuitive that the point at which one can no longer infer a WGD is when more than half of all genes duplicated have returned to single-copy. As there are only ~550 homoeologs remaining in the *S*. *cerevisiae* genome (Byrne and Wolfe 2005), it may therefore be surprising that we correctly favor allopolyploidy over no polyploidy in our analysis of yeast genomes. The key factor in our statistical power to reject a history without WGD appears to be the fact that there are distinct homoeologs lost or preserved in different lineages (cf. Scannell et al. 2007). It is the sum total of these homoeologs that enable us to infer an allopolyploid history, such that the large number of polyploid species included in our analysis has helped to support this inference. We caution users of our method that any conclusions concerning the presence or absence of polyploidy is therefore dependent on many factors, including the age of the polyploidy event, the rate of duplicate gene loss (and the asymmetry of loss between homoeologs), and the number of polyploid lineages sampled.

Importantly, errors in gene tree reconstruction or incongruence caused by biological phenomena such as incomplete lineage sorting are problems in any tree-based analysis of polyploidy, and methods to both quantify and account for them have been established in other contexts (e.g. Jones et al. 2013; Zwickl et al. 2014). Gene tree incongruence is also a problem for reconciliation algorithms (Hahn 2007), though solutions have been proposed to deal with incongruence due to ILS (e.g. Vernot et al. 2008; Rasmussen and Kellis 2012; Yu et al. 2013). In the future it will be valuable to implement a similar solution for the case of reconciliation applied to WGDs, or a solution based on ILS in species networks (Jones et al. 2013). Methods that do not rely on gene tree reconstruction are also viable alternatives for very recent polyploidy events (Roux and Pannell 2015).

The algorithm and associated software presented here should allow researchers to re-examine many published cases of polyploidy, in order to determine whether these events were auto- or allopolyploidy. While many clades of plants often have multiple WGD events within them, our re-analysis of the wheat data gives us confidence that our method can be expanded to identify multiple polyploidy events in the same tree. For cases with only a single WGD, our method provides accounting of duplication and loss, as well as the placement of these events in a phylogenetic context.

## Acknowledgements

We thank Clara Boothby for comments on the manuscript, Marina Marcet-Houben for helpful feedback regarding the yeast data, Ben Moore for information about the wheat data, and Luay Nakhleh and Celine Scornavacca for pointing out important missing references. Thomas Marcussen, one anonymous reviewer, and Susanne Renner also provided helpful comments. This work was funded by National Science Foundation grant DBI-1564611 to MWH.

---

**Figure S1: Scenarios with various levels of extinction of parental lineages.** In each panel, A’ and B’ have hybridized to form the polyploid clade, P. We define A’ and B’ as the parental *species.* A is more closely related to A’ than B’, so we say that A and A’ make up one parental *lineage* of the polyploid (teal). Likewise, B is more closely related to B’ than to A’, thus making a second parental *lineage* (purple). *(a)* With no extinctions and all parental species sampled, GRAMPA will recover the correct, allopolyploid MUL-tree and correctly infer the parental lineages. *(b)* If a single parental *species* is extinct or unsampled (A’ in this instance), but other species from that parental *lineage* are extant and sampled, GRAMPA will still infer an allopolyploid MUL-tree and correctly identify the parental lineages. This would still be the case if both parental *species* were extinct. *(c)* Even if an entire parental *lineage* is extinct or unsampled, GRAMPA will still infer allopolyploidy and correctly imply the second parental lineage. Only one species from one parental lineage need be present to be able to correctly infer allopolyploidy. *(d)* In cases where both parental lineages of an allopolyploid are entirely extinct, GRAMPA will incorrectly infer autopolyploidy with a WGD occurring between the two sub-genomes of the polyploid clade (orange circle in the inferred network).

**Figure S2: An example scenario of GRAMPA’s heuristic speed-ups.** Given the MUL-tree in (a) with two sub-genomes represented in the XYZ clade, the base algorithm will try all possible combinations of initial maps for genes from polyploid species, as in (b). If polyploid species group together in the gene tree, the most parsimonious initialization will always have them mapping to the same sub-genome. We collapse these into groups to reduce the number of possible maps we have to try, as in (c). Furthermore, if the polyploid lineage has a sister clade in the singly-labeled species phylogeny (species A, in this case), we can look for the same pattern in the gene trees and fix the maps of any groups we find, as in (d).

**Figure S3: The singly-labeled species tree topology used to generate MUL-trees and gene trees for the manual reconciliations and an example MUL-tree.** *(a)* Node H1 defines the polyploid clade made up of species X, Y, and Z. The node ancestral to species C and D is specified as H2 as an example, however any other node outside of the polyploid clade can be considered as H2. If H1 and H2 are the same node, the resulting MUL-tree is indicative of autopolyploidy. *(b)* The MUL-tree generated by using the H1 and H2 nodes defined in *(a).*

**Figure S4: The various simulation scenarios used to test GRAMPA.** *(a)* Gene trees simulated with allopolyploidy. The allopolyploid genome topology (G) was used as input for JPrIME. Three scenarios of ILS were considered, with the two discordant topologies also being used as input to JPrIME. Additionally, five scenarios of various gene gain-loss (G-L) rates were considered. This led to 15 sets of 1000 gene trees that were subsequently used as inputs to GRAMPA, along with both singly-labeled topologies. *(b)* Gene trees simulated with autopolyploidy. The autopolyploid genome topology G was used as input for JPrIME with varying scenarios of ILS and G-L. *(c)* Gene trees simulated with no polyploidy. The species topology (S) was used as input to JPrIME with varying scenarios of ILS and G-L.

**Figure S5: Yeast phylogeny.** (***a***) The phylogeny from Marcet-Houben and Gabaldón (2015) of 27 yeast species used in the analysis. The putative H1 node is n5 while the inferred H2 node is n3. (**b**) A simplified version of the species tree showing only the clades and nodes of interest.

**Figure S6: Wheat phylogeny**. (*a*) The full species tree used for the bread wheat data. Common bread wheat, *Triticum aestivum* is represented by three sub-genomes, A, B, and D. Nine other closely related species were considered. (**b)** A simplified version of the tree showing only the three sub-genomes of bread wheat (labeled A, B, and D), and the donor species of two of the sub-genomes, *A. tauschii* and *T. urartu* (abbreviated AT and TU, respectively).

**Figure S7: Demonstration of penalties when reconciling polyploids to singly-labeled trees.** *(a)* When allopolyploidy has occurred, as shown in the MUL-tree on the left, there is a much higher score for reconciling to a singly-labeled tree when both homoeologous copies are retained (gene tree 1) or when the copy that is not represented in the singly-labeled tree is retained while the other is lost (gene tree 3), as indicated by the negative difference in reconciliation scores (AS). We see only relatively small penalties for reconciling to the correct MUL-tree in scenarios that favor the singly-labeled tree, such as retention of the homoeologous copy that is represented in the singly-labeled tree and loss of the other copy (gene trees 2 and 4). ***(b)*** When autopolyploidy has occurred, as shown in the MUL-tree on the left, the penalties for reconciling to a singly-labeled tree still exist when both ohnologs are present, but are much smaller (gene tree 1). In fact, since there is only one placement of both sub-genomes, a loss of either ohnolog favors the singly-labeled tree. This means we can still distinguish autopolyploidy from no polyploidy, but have much less power to do so than with allopolyploidy (i.e. many more ohnologs must be retained to reject no polyploidy).

**Figure S8: Wheat polyploidy events inferred by GRAMPA when considering only 2 sub-genomes.** In each panel, the top tree is the species tree input for GRAMPA and the bottom panel is the optimal MUL-tree inferred. The bread wheat species, *Triticum aestivum,* is abbreviated TA. (**a**) Only sub-genomes A and D are present in the gene trees and the position of TA in the input singly-labeled tree is at sub-genome A. The optimal MUL-tree shows the correct placement of sub-genome D. (**b**) Only sub-genomes A and B are present in the gene trees and the position of TA in the input singly-labeled tree is at sub-genome A. The optimal MUL-tree shows the correct placement of sub-genome B. (**c**) Only sub-genomes D and B are present in the gene trees and the position of TA in the input singly-labeled tree is at sub-genome D. The optimal MUL-tree shows the correct placement of sub-genome B.

**Figure S9: Wheat polyploidy events inferred by GRAMPA when considering all 3 sub-genomes.** The top two scoring MUL-trees returned by GRAMPA when considering all three *T. aestivum* (TA) sub-genomes. **(*a*)** The input species tree given to GRAMPA with only sub-genome A represented. **(b)** The best scoring MUL-tree finds sub-genome B as the other TA lineage. **(d)** The second best scoring MUL-tree finds sub-genome D as the other TA lineage.

## References

Adams KL, Wendel JF. 2005. Polyploidy and genome evolution in plants. Curr Opin Plant Biol. 8(2):135–141.

Altenhoff AM, Škunca N, Glover N, Train C-M, Sueki A, Piližota I, Gori K, Tomiczek B, Müller S, Redestig H, Gonnet GH, Dessimoz C. 2015. The OMA orthology database in 2015: Function predictions, better plant support, synteny view, and other improvement. Nucleic Acids Res. 43(Database issue):D240–D249.

Amores A, Force A, Yan YL, Joly L, Amemiya C, Fritz A, Ho RK, Langeland J, Prince V, Wang YL, Westerfield M, Ekker M, Postlethwait JH. 1998. Zebrafish hox clusters and vertebrate genome evolution. Science. 282(5394): 1711–1714.

Barker MS, Kane NC, Matvienko M, Kozik A, Michelmore RW, Knapp SJ, Rieseberg LH. 2008. Multiple paleopolyploidizations during the evolution of the Compositae reveal parallel patterns of duplicate gene retention after millions of years. Mol Biol Evol. 25(11):2445–2455.

Barker MS, Vogel H, Schranz ME. 2009. Paleopolyploidy in the Brassicales: analyses of the *Cleome* transcriptome elucidate the history of genome duplications in *Arabidopsis* and other Brassicales. Genome Biol Evol. 1:391–399.

Barker MS, Arrigo N, Baniaga AE, Li Z, Levin DA. 2015. On the relative abundance of autopolyploids and allopolyploids. New Phytol. 210:391–398.

Blanc G, Hokamp K, Wolfe KH. 2003. A recent polyploidy superimposed on older large-scale duplications in the *Arabidopsis* genome. Genome Res. 13(2):137–144.

Blanc G, Wolfe KH. 2004. Widespread paleopolyploidy in model plant species inferred from age distributions of duplicate genes. The Plant Cell.; 16(7):1667–1678.

Bolser DM, Kerhornou A, Walts B, Kersey P. 2015. Triticeae resources in Ensembl Plants. Plant Cell Physiol. 56(1):e3.

Bowers JE, Chapman BA, Rong J, Paterson AH. 2003. Unravelling angiosperm genome evolution by phylogenetic analysis of chromosomal duplication events. Nature. 422(6930): 433–438.

Brassac J, Blattner FR. 2015. Species-level phylogeny and polyploid relationships in *Hordeum* (Poeaceae) inferred by next-generation sequencing and *in silico* cloning of multiple nuclear loci. Syst Biol. 64(5):792–808.

Byrne KP, Wolfe KH. 2005. The Yeast Gene Order Browser: Combining curatedhomology and syntenic context reveals gene fate in polyploid species. Genome Res. 15(10): 1456–1461.

Cannon SB, McKain MR, Harkess A, Nelson MN, Dash S, Deyholos MK, Peng Y, Joyce B, Stewart CN Jr, Rolf M, Kutchan T, Tan X, Chen C, Zhang Y, Carpenter E, Wong GK, Doyle JJ, Leebens-Mack J. 2015. Multiple polyploidy events in the early radiation of nodulating and nonnodulating legumes. Mol Biol Evol. 32(1): 193–210.

Chalup L, Grabiele M, Soís Neffa V, Seíjo G. 2012. Structural karyotypic variability and polyploidy in natural populations of the South American Lathyrus nervosus Lam. (Fabaceae). Plant Syst Evol. 298:761.

Crow KD, Wagner GP. 2006. What is the role of genome duplication in the evolution of complexity and diversity? Mol Biol Evol. 23(5):887–892.

Cui L, Wall PK, Leebens-Mack JH, Lindsay BG, Soltis DE, Doyle JJ, Soltis PS, Carlson JE, Arumuganathan K, Barakat A, Albert VA, Ma H, dePamphipilis CW. 2006. Widespread genome duplications throughout the history of flowering plants. Genome Res. 16(6):738–749.

Dehal P, Boore JL. 2005. Two rounds of whole genome duplication in the ancestral vertebrate. PLoS Biol. 3(10):e314.

Durand D, Halldórsson BV, Vernot B. 2006. A hybrid micro-macroevolutionary approach to gene tree reconstruction. J Comput Biol. 13(2):320–335.

Edger PP, Heidel-Fischer HM, Bekaert M, Rota J, Glöckner G, Platts AE, Heckel DG, Der JP, Wafula EK, Tang M, Hofberger JA, Smothson A, Hall JC, Blanchette M, Bureau TE, Wright SI, dePamphilis CW, Schranz ME, Barker MS, Conant GC, Wahlberg N, Vogel H, Pires JV, Wheat CW. 2015. The butterfly plant arms-race escalated by gene and genome duplications. Proc Natl Acad Sci USA. 112(27):8362–8366.

Fitch WM. 1970. Distinguishing homologous from analogous proteins. Syst Zool. 119(2): 99–113.

Glover NM, Redestig H, Dessimoz C. 2016. Homoeologs: What are they and how do we infer them? Trends Plant Sci. S1360–1385(16):00059–5.

Goodman M, Czelusniak J, William Moore G, Romero-Herrera AE, Matsuda G. 1979. Fitting the gene lineage into its species lineage, a parsimony strategy illustrated by cladograms constructed from globin sequences. Syst Biol. 28(2):132–163.

Guigó R, Muchnik I, Smith TF. 1996. Reconstruction of ancient molecular phylogeny. Mol Phylogenet Evol. 6(2): 189–213.

Hahn MW. 2007. Bias in phylogenetic tree reconciliation methods: implications for vertebrate genome evolution. Genome Biol. 8(7):R141.

Huber KT, Moultan V. 2006. Phylogenetic networks from multi-labelled trees. J Math Biol. 52(5):613–632.

Huber KT, Oxelman B, Lott M, Moultan V. 2006. Reconstructing the evolutionary history of polyploids from multilabeled trees. Mol Biol Evol. 23(9):1784–1791.

Huson DH, Rupp R, Scornavacca C. 2006. Phylogenetic networks from trees. Phylogenetic Networks, Cambridge University Press.

Jiao Y, Wickett NJ, Ayyampalayam S, Chanderbail AS, Landherr L, Ralph PE, Tomsho LP, Hu Y, Liang H, Soltis PS, SOltis DE, Clifton SW, Schlarbaum SE, Schuster SC, Ma H, Leebns-Mack J, dePamphilis CW. 2011. Ancestral polyploidy in seed plants and angiosperms. Nature. 473(7345):97–100.

Jones G, Sagitov S, Oxelman B. 2013. Statistical inference of allopolyploid species networks in the presence of incomplete lineage sorting. Syst Biol. 62(3):467–478.

Junier T, Zdobnov EM. 2010. The Newick Utilities: high-throughput phylogenetic tree processing in the UNIX shell. Bioinformatics. 26:1669–1670.

Kellis M, Birren BW, Lander ES. 2004. Proof and evolutionary analysis of ancient genome duplication in the yeast *Saccharomyces cerevisiae*. Nature. 428(6983):617–624.

Kersey PJ, Allen JE, Armean I, Boddu S, Bolt BJ, Carvalho-Silva D, Christensen M, Davis P, Falin LJ, Grabmueller C, Humphrey J, Kerhornou A, Khobova J, Aranganathan NK, Langridge N, Lowy E, McDowall MD, Maheswari U, Nuhn M, Kee Ong C, Overduin B, Paulini M, Pedro H, Perry E, Spudich G, Tapanari E, Walts B, Williams G, Tello-Ruiz M, Stein J, Wei S, Ware D, Bolser DM, Howe KL, Kulesha E, Lawson D, Maslen G, Staines DM. 2016. Ensembl Genomes 2016: more genomes, more complexity. Nuc Acids Res. 44(D1):D574–D580.

Kim C, Wang X, Lee TH, Jakob K, Lee GJ, Paterson AH. 2014. Comparative analysis of *Miscanthus* and *Saccharum* reveals a shared whole-genome duplication but different evolutionary fates. Plant Cell. 26(6):2420–2429.

Li Z, Baniaga AE, Sessa EB, Scascitelli M, Graham SW, Rieseberg LH, Barker MS. 2015. Early genome duplications in conifers and other seed plants. Sci Adv. 1(10):e1501084.

Linder CR, Rieseberg LH. 2004. Reconstructing patterns of reticulate evolution in plants. Am J Bot. 91(10):1700–1708.

Lott M, Spillner A, Huber KT, Petri A, Oxelman B, Moulton V. 2009. Inferring polyploid phylogenies from multiply-labeled gene trees. BMC Evol Biol. 9:216.

Lynch M, Conery JS. 2000. The evolutionary fate and consequences of duplicate genes. Science. 290(5495): 1151–1155.

Lynch M, Force A. 2000. The probability of duplicate gene preservation by subfunctionalization. Genetics. 154(1):459–473.

Mandáková T, Joly S, Krzywinski M, Mummenhoff K, Lysak MA. 2010. Fast diploidization in close mesopolyploid relatives of *Arabidopsis*. Plant Cell. 22(7):2277–2290.

Marcussen T, Jakobsen KS, Danihelka J, Ballard HE, Blaxland K, Brysting AK, Oxelman B. 2012. Inferring species networks from gene trees in high-polyploid North American and Hawaiian violets (*Viola*, Violaceae). Syst Biol. 61(1): 107–126.

Marcussen T, Sandve SR, Heier L, Spannagl M, Pfeifer M, International Wheat Genome Sequencing Consortium, Jakobsen KS, Wulff BB, Steuernagel B, Mayer KF, Olsen OA. 2014. Ancient hybridizations among the ancestral genomes of bread wheat. Science. 345(6194):1250092.

Marcussen T, Heier L, Brysting AK, Oxelman B, Jakobsen KS. 2015. From gene trees to a dated allopolyploid network: insights from the angiosperm genus *Viola* (Violaceae). Syst Biol. 64(1):84–101.

Marcet-Houben M, Gabaldón T. 2015. Beyond the whole genome duplication: Phylogenetic evidence for an ancient interspecies hybridization in the baker’s yeast lineage. PLoS Biol. 13(8):e1002220.

Mayrose I, Zhan SH, Rothfels CJ, Magnuson-Ford K, Barker MS, Rieseberg LH, Otto SP. 2011. Recently formed polyploid plants diversify at lower rates. Science. 333:1257.

Muir CD, Hahn MW. 2015. The limited contribution of reciprocal gene loss to increased speciation rates following whole genome duplication. The American Naturalist. 185:70–86.

Ohno S. 1970. Evolution by gene duplication. Springer-Verlag

Page RDM. 1994. Maps between trees and cladistic analysis of historical associations among genes, organisms, and areas. Syst Biol. 43(1):58–77.

Petersen G, Seberg O, Yde M, Berthelsen K. 2006. Phylogenetic relationship of *Triticum* and *Aegilops* and evidence for the origin of the A, B, and D genomes of common wheat (*Triticum aestivum*). Mol Phylogenet Evol. 39(1):70–82.

Popp M, Oxelman B. 2001. Inferring the history of the polyploid *Silene aegaea* (Caryophyllaceae) using plastid and homoeologous nuclear DNA sequences. Mol Phylogenet Evol. 20(3):474–481.

Rabier C-E, Ta T, Ané C. 2014. Detecting and locating whole genome duplications on a phylogeny: A probabilistic approach. Mol Biol Evol. 31(3):750–762.

Rasmussen MD, Kellis M. 2012. Unified model of gene duplication, loss, and coalescence using a locus tree. Genome Res. 22(4):755–765.

Roux C, Pannell JR. 2015. Inferring the mode of origin of polyploidy species from next-generation sequence data. Mol Ecol. 24:1047–1059.

Scannell DR, Frank AC, Conant GC, Byrne KP, Woolfit M, Wolfe KH. 2007. Independent sorting-out of thousands of duplicated gene pairs in two yeast species descended from a whole-genome duplication. Proc Natl Acad Sci USA. 104(20):8397–402.

Sjöstrand J, Arvestad L, Lagergren J, Sennblad B. 2013. GenPhyloData: Realistic simulation of gene family evolution. BMC Bioinformatics. 14:209.

Soltis PS, Soltis DE. 2009. The role of hybridization in plant speciation. Annu Rev Plant Biol. 60:561–588.

Tiley GP, Ané C, Burleigh JG. 2016 Evaluating and characterizing ancient whole-genome duplications in plants with gene count data. Genome Biol Evol. 8(4): 1023–1037.

To TH, Scornavacca C. 2015. Efficient algorithms for reconciling gene trees and species networks via duplication and loss events. BMC Genomics. 16(10):S6.

Van de Peer Y. 2004. Computational approaches to unveiling ancient genome duplications. Nat Rev Genet. 5(10):752–763.

Vernot B, Stolzer M, Goldman A, Durand D. 2008. Reconciliation with non-binary species trees. J Comput Biol. 15(8):981–1006.

Werth CR, Windham MD. 1991. A model for divergent, allopatric speciation of polyploid Pteridophytes resulting from silencing of duplicate-gene expression. The American Naturalist. 137(4):515–526.

Wolfe KH. 2000. Robustness-it’s not where you think it is. Nat Genet. 25(1):3–4.

Wolfe KH, Shields DC. 2007. Molecular evidence for an ancient duplication of the entire yeast genome. Nature. 287(6634):708–713.

Yu Y, Barnett RM, Nakhleh L. 2013. Parsimonious inference of hybridization in the presence of incomplete lineage sorting. Syst Biol. 62(5):738–751.

Zmasek CM, Eddy SR. 2001. A simple algorithm to infer gene duplication and speciation events on a gene tree. Bioinformatics. 17(9):821–828.

Zwickl DJ, Stein JC, Wing RA, Ware D, Sanderson MJ. 2014. Disentangling methodological and biological sources of gene tree discordanc on *Oryza* (Poaceae) chromosome 3. Syst Biol. 63(5):645–659.

